# Unraveling the aggregation dynamics of amyloidogenesis: A data-driven mathematical modeling approach

**DOI:** 10.1101/2020.01.06.895649

**Authors:** Baishakhi Tikader, Samir K. Maji, Sandip Kar

**Affiliations:** Department of Chemistry, IIT Bombay, Powai, Mumbai - 400076, India; Department of Biosciences and Bioengineering, IIT Bombay, Powai, Mumbai - 400076, India

## Abstract

Proteins often get misfolded into highly ordered amyloid aggregates that are found to be associated with various neurodegenerative diseases. However, the dynamical events that lead to such self organization of different types of proteins are still poorly understood due to the experimentally untractable complex molecular mechanism, which governs the process of amyloidogenesis. Herein, we propose a generic and data-driven mathematical modeling approach that enables us to decipher a most probable molecular mechanism responsible for amyloidogensis in a context dependent manner. The preferred model efficiently elucidates various aspects of amyloid forming kinetics as a function of initial protein concentration, and qualitatively predicts the dynamics of amyloidogenesis for point mutated proteins. Importantly, the model analysis reveals that intermediate aggregates formed just after the primary necleation steps plays a vital role in dictating the nature of amyloid forming kinetics. Moreover, our model makes experimentally feasible insightful predictions to formulate better therapeutic measures in future to counter unwanted amyloidogenesis by just fine-tuning the molecular interactions related to amyloidogenesis.

**Author Summary:** In cells, proteins mostly function by being in a native folded conformation. However, under certain circumstances, some of these proteins undergo high degree of aggregation and eventually misfold to form highly stable amyloid aggregates that are often involved in different neurodegenerative disease progression and pathogenesis. To develop appropriate therapeutic strategies to resist such oligomerazation of protein, it is imperative to unravel the molecular interactions that essentially organize the dynamical events of amyloidogenesis. We have proposed a generic computational modeling framework to elucidate the important molecular events that govern the dynamics of amyloid formation. Our model reconciles diverse set of experimental observations including different mutant phenotyes from a dynamical perspective, and predicts possible ways to control the kinetics of amyloid formation experimentally. In future, these insights will be helpful to design drugs to treat patients with neurodegenerative disorders. Importantly, our model, in general, will be quite useful to figure out the important molecular events that are orchestrating the amyloidogenesis of different amyloid forming proteins.

## Introduction

Biological function of a protein is intricately related to its structural state ([1]). Under normal circumstances, a protein functions by maintaining its stable natively folded structure. Intriguingly, often proteins misfold to produce thermodynamically highly stable aggregated states (commonly known as amyloid fibrils), which under different physiological contexts are associated with a number of diseases states such as neurodegenerative disease (like Alzheimer’s, Parkinson’s and Huntington’s diseases), Type-II diabetes and prion disease ([2],[3],[4],[5],[6]). At the same time, higher stability and diverse functional activities of different amyloid fibrils can in principle be utilized to fabricate novel biomaterials for various application purposes. However, the aggregation mechanisms and the related dynamical transitions that result in amyloid fibril formation in different scenarios are extremely complex and unpredictable in nature ([7],[8]). The conventional experimental studies related to aggregation dynamics of proteins suffer from lack of mechanistic insight, and getting a quantitative account of the dynamics of amyloid formation is still hard to achieve. Recently, attempts have been made in addressing these issues at the single molecule level ([9],[10],[11]), however, a comprehensive quantitative understanding about the amyloid fibril formation dynamics still mostly remains elusive. Nonetheless, it is evident that such an effort is imperative to develop methods to fine-tune the dynamics of amyloid fibril formation in a context dependent manner.

Experimentally, it has been observed that the amyloid fibril formation normally begins with primary nucleation step, where monomers of the subsequent protein initiate the aggregation process ([12]). This step is followed by several other processes such as elongation, secondary nucleation/fragmentation, and even some conformation changes to finally produce the amyloid fibril ([8]). However, it is experimentally highly challenging to disentangle distinctively how each these phases are influenced by the different molecular processes involved in amyloid fibril growth. In this context, a number of theoretical models have been developed to explain the complicated and heterogeneous molecular mechanism involved in aggregation process. Researchers have employed the concept of nucleation dependent polymerization (NDP) to explain the various aspects (like sigmoidal growth kinetics, effect of seeding on lag time, fibril elongation etc.) of amyloid formation kinetics qualitatively ([8],[13],[14],[15],[16],[17],[18]). Several other views such as existence of a super critical concentration controlling the NDP process, and fibril positively regulating the fibrillation has been proposed to investigate the kinetics of amyloidogenesis ([19],[20]). However, in these approaches, the possibilities of conformational transitions have been totally overlooked.

On the other hand, Lumry et al. hypothesize that the amyloidogenesis is mainly driven by folding-unfolding structural transition but mostly they disregarded the sigmoidal kinetic features of fibril formation ([20],[21],[22],[23]). Recently, it has been suggested that the existence of certain intermediate has been identified during the transition from native structure to mature fibril formation process ([24],[25],[26],[27]). This clearly indicates that a complex set of mechanisms and pathways are involved in developing the amyloid fibril. Additionally, the dynamics of amyloidogenesis differs considerably depending on several experimental parameters (for example; initial total protein concentration, point mutations etc.). Not only that, the observed quantities (for example; the lag time) vary quite significantly depending on how the experiments are performed to quantify the kinetic information of amyloid formation. Thus, the question remains, how to tackle such kind of complex and heterogeneous experimental data to get an improved and holistic view of dynamical transitions pertinent to amyloid fibril formation?

In literature, mathematical and computational modeling studies played pivotal role to gain crucial insight related to various biological processes underlying the aggregation kinetics for different proteins under varied experimental conditions. These models have evolved with time considerably, however, each of these mathematical and computational approaches is found to be limited in one way or other. The molecular dynamics approach to resolve the dynamics of fibril formation can only provide the dynamical information for about tens (10’s) of microseconds and cannot handle a large pull of molecules. Consequently, it is difficult to reconcile various experimental results, where the numbers of molecules are large and the process of fibrillation takes a much longer time. Coarse graining such systems show a bit of improvement but lacks the molecular interpretations of the underlying mechanism of fibril formation dynamics. Yet, very few models considered all the mechanistic details of amyloid fibril formation, and mostly provided a qualitative description of the overall process. Thus, predictive power of those models is limited.

Herein, we have employed a mathematical modeling approach to construct a generic model comprising most of the mechanistic complexity, which can provide inclusive insight about various microscopic events that involved in the aggregation process. We have proposed a data-driven mathematical model and calibrate our model with experimental data set and try to give a more global picture of experimentally observed kinetic behavior in case of any amyloid forming protein. Consequently, we have implemented a systematic testing of different possible assembly network and found a very important feature that the linear dependence of secondary nucleation rate on the elongation rate favors the aggregation process (**S2 Text**). Importantly, from a dynamical perspective, our model explains; (i) the quantitative variations of the kinetics of amyloid fibril formation observed with Thioflavin T (ThT) assay in a concentration dependent manner, (ii) the diverse kinetic behavior of any mutants of a specific amyloid forming protein, (iii) kinetics of seeding experiments performed by adding elongation aggregated structures formed during amyloidogenesis, and (iv) even predicts how the amyloid forming kinetics can be altered by perturbing different steps responsible for amyloid formation. Moreover, the proposed dynamical model due to its comprehensive nature, gives a better description of the fibrillation process, postulates novel mechanistic hypotheses, and provides further insights to fine-tune the fibrillation dynamics in a preferred way for both therapeutic and other application purposes.

## The Model

In last few decades, mathematical models are often employed to understand the dynamics of amyloid fibril formation. Keeping this in mind, our aim to propose a more generic and informative model of amyloid fibril formation that can be applicable to any protein, which has a propensity to form amyloid like aggregates. Our model is not only applicable for intrinsically disordered protein ([28]) (like *α*-Synuclein), where an unstructured protein transits from random coil like state to intermediate elongation unit, which eventually converts to fibril like state (three state transition, RC → intermediate → *β*-sheet) ([24],[25],[26],[27]), but the model also holds good for all proteins, which shows only random coil state to fibril like state transition without going through any other intermediate conformations ([29],[30]) (two state transition, RC → *β*-sheet).

In our proposed model (**Fig. 1** and **Figure S1**), we have incorporated most of the mechanistic details that are proposed till date in literature to obtain a more universal picture of amyloidogenesis. Therefore, our model involves a number of species, and we have described each model variable representing these species in **Table S1**.

**Fig. 1.**
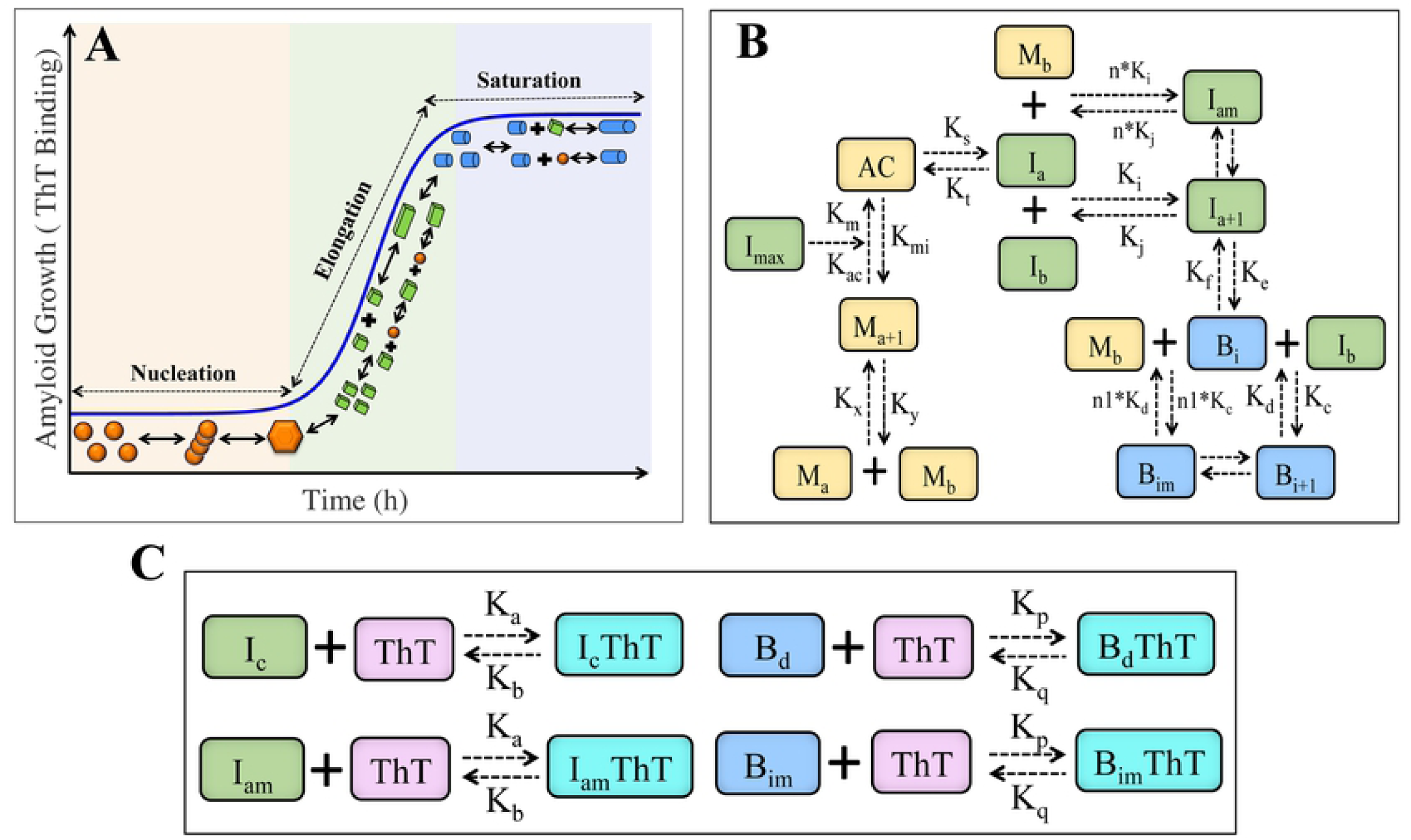
The proposed interaction network of protein aggregation describing the dynamics of amyloidogenesis. **(A)** Schematic representation of different microscopic molecular mechanism that occurs at the different phases of the aggregation process of any amyloid forming protein. **(B)** Proposed protein interaction network for aggregation process that leads to amyloid formation for different proteins. Various indices (like a = (1-3), b = 1, i = (1-2), m = (a-c), c = (1-4), d = (1-3)) are used to indicate the polymeric forms of different species such as M, I, B etc. The model includes the concepts of; (i) primary nucleation process, where random coil states (*M*_*a*_) self-associates to form partially folded conformation (we termed activated complex, AC) via (ii) normal or facilitated molecular rearrangements from random coil state (for example; *M*_4_→*AC* transition), (iii) This activated complex then undergoes conformational transition (for example; *AC*→*I*_1_) to form elongated building block or fibrillar building block such as *I*_*c*_ *or B*_*d*_, which then either (iv) self-associate (for example; elongation process of basic elongation units (*I*_*c*_) and basic fibril forming units *B*_*d*_), and/or (v) elongated in presence of monomer (secondary nucleation) to form mature fibril unit (for example, *B*_*d*_). **(C)** Interactions network representing the association and dissociation kinetics of Thioflavin T (ThT) with elongation units (I_c_/I_am_) and fibril formation units (B_d_/B_im_) of the amyloid forming protein. Details about the network have been described in **S1 Text** of Supplementary information.

We began constructing our model by considering the nucleation-dependent polymerization phenomena, as it is one of the most crucial steps for amyloidogenesis ([31],[32],[33]). We assumed that the monomers (M_1_) can self-assemble to each other to form a nucleus like structure, from which the basic fibril elongation process can initiate. It is important to mention here that in most of the kinetic models proposed earlier, the structural transition of nucleus to reach an activated like state has been ignored, though conformal changes during self-assemble process is an essential process during amyloid formation ([22],[23]). In our model, we have hypothesized that the nucleus like state (M_4_) can undergo conformational rearrangement to give an activated state (AC), which can act as a seed for the continuation of the further processes. In literature, it has also been reported that natively unstructured proteins may undergo; (i) large structural transition from random coil to β-sheet rich fibril directly ([34],[35]), or (ii) the transition can happen via some partially folded intermediate (such as helix rich intermediate) ([24],[25],[26],[27],[34]). Under such circumstances, the oligomeric unit I_1_ can be interpreted either as the basic minimum elongation unit of fibril (for the case (i)), or as the basic minimum helix rich intermediate unit (for the case (ii)). These elongation units can further self-associate to form the seed for next structural transition (for the case (ii)), or the higher ordered stable form of mature fibrils (for the case (i)), which again can grow via self-association or by secondary nucleation to finally form mature fibril (for details see **S1 Text**, and the related discussion therein). Thus, our modeling framework can be applied to any protein that under certain condition leads to amyloid formation.

We have translated all these interactions (**Fig. 1** and **Fig S1**) into sets of ordinary differential equations (**Table S2**). Importantly, our model is completely based on mass-action kinetic terms (**Table S2**), and depicts the abovementioned interactions in a highly simplified manner. The model further involves few algebraic relations (**Table S3**) for ease of calculation. Additionally, the model consists of a number of unknown kinetic parameters (**Table S4**) representing the interaction network described in **Fig. 1**. It is extremely important to get a reasonable estimate of these kinetic parameters involved in the proposed model to elucidate and investigate the inherent mechanism behind this heterogeneous process. In our case, we focused on the kinetic data reported by Ghosh *et al*. related to *α*-Synuclein protein aggregation ([12]) to have a preliminary estimate of the kinetic parameters involved in our model. Our principle purpose here is to obtain a comprehensive understanding of the kinetics of amyloid formation corroborating with various experimental observations related to *α*-Synuclein protein and other amyloid forming proteins, and to expand these insights to dissect the underlying dynamics of amyloidogenesis.

## Results and discussion

### Model reproduces the concentration dependency of the aggregation kinetics

Experimentally, Ghosh et al. had simultaneously quantified the kinetics of amyloidogenesis by performing experiments with Thioflavin T, and measured the concentration of the total intermediate (*I*_*total*_) with time during the course of amyloid formation of *α*-Synuclein protein ([12])

To begin with, by considering these two types of experimental data sets, we separately optimized different ensemble of models that either represent the interaction network described in **Fig. 1** or it’s different variants (**Table S5, Fig S1**) to come up with an optimal network structure that will best fit the experimental data under consideration. The best network model (Model 1A, **Table S5**) was selected on the basis of statistical (*χ*^2^ values and Akaike information criteria (AIC)) criteria (for details see **Method** section and **S2 Text**). In **Table S4**, we have given the parameter set obtained for the best fit after fitting the model (Model 1A, **Table S5**) with the experimental data shown in **Fig. 2A-B**. This depicts that our proposed model can adequately elucidate all the important features of the sigmoidal amyloid growth kinetics observed (**Fig. 2A-B**) experimentally for 300 *μ*M total protein concentration. Further, we have obtained additional insights about the underlying network structure that eventually leads to such kind of kinetics.

**Fig. 2.**
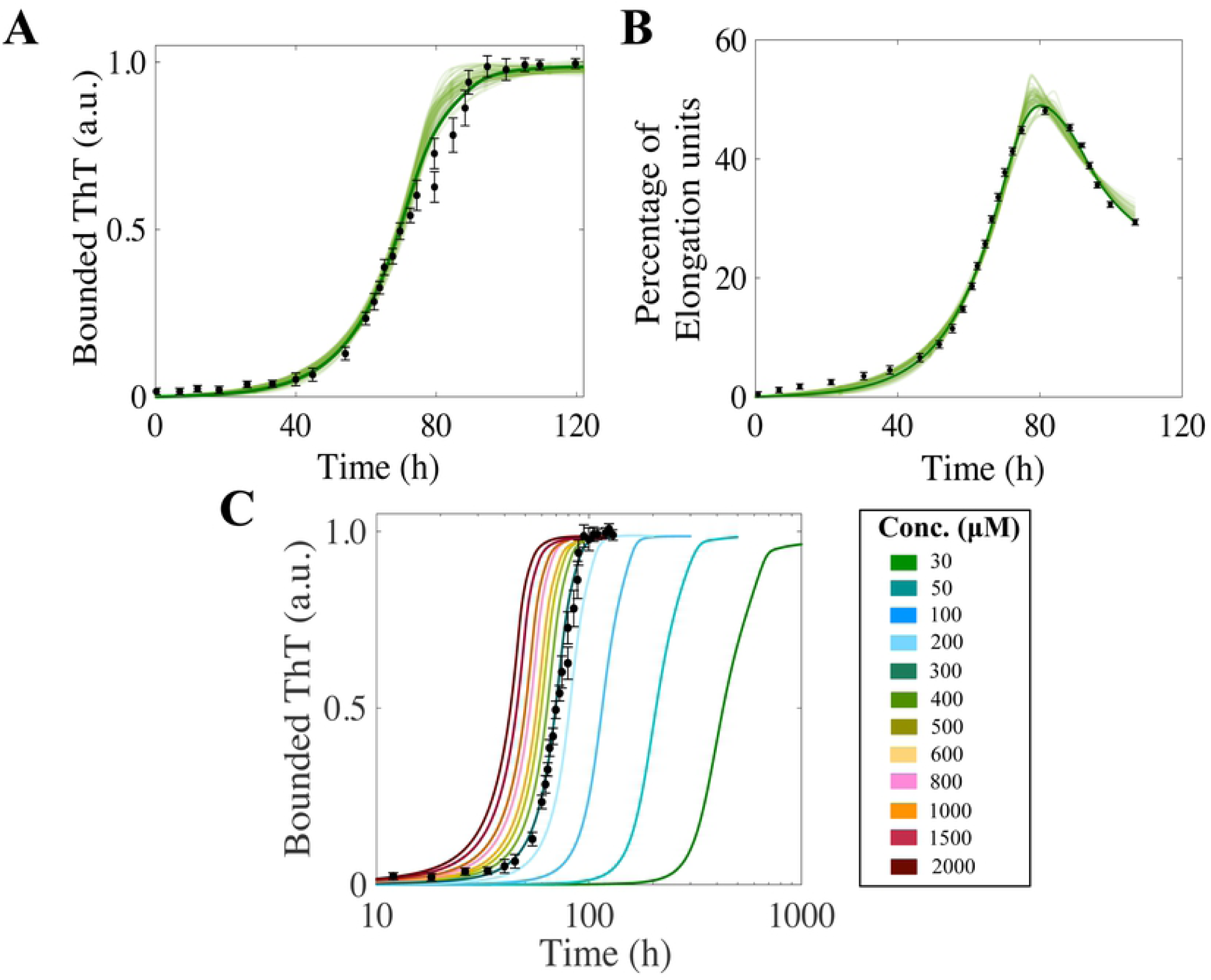
Model reconciles the concentration dependency of amyloid forming kinetics. **(A)** The time courses (best 50 fits are shown, where the solid green line represents the best-fitted trajectory) of the total ThT bounded aggregates (**S2 Text**) obtained by fitting the experimental data (black dots with error bars, for 300 *μ*M total protein concentration). **(B)** The time courses (best 50 fits are shown, where the solid green line represents the best-fitted trajectory) of % of total intermediate aggregates obtained by fitting the experimental data (black dots with error bars). **(C)** The model predicted time courses of total ThT bounded aggregates under different total protein concentrations on a logarithm scale. Here, we only varied the total protein concentration by keeping all other parameters same as given in **Table S4**.

Interestingly, in literature, it has been reported that the kinetics of amyloidogenesis changes its nature as a function of initial protein concentration ([12],[36]). We asked whether our model could predict the experimentally observed kinetics for the total protein concentration ranging from low to intermediate to high values? Can it provide up to what amount of total protein concentration, the sigmoidal growth curve could be maintained? To investigate these important questions, in **Fig. 2C**, we have demonstrated the model predicted time profiles (with best-fitted parameters given in **Table S4**) of the total ThT bounded aggregates as a function of time. Our model predictions reconciled the experimentally observed kinetics, when the total protein concentrations are either decreased or increased systematically. Our model is extremely successful in this perspective producing the consequent kinetics at specific conditions and maintains a sustain feature of sigmoidal growth kinetics with no perceptible changes in the slope of the sigmoidal growth, though we have fitted our model only at a particular concentration. The subsequent increase or decrease in the total protein concentration essentially speeds up or slows down the self-assembly process in every elementary step of aggregation, as a result either the entire rise up or the overall drop in the sigmoidal growth kinetics has been observed that has been flowed in the bounded ThT profiles. This reflects that our model can successfully generates the amyloid kinetics for low to intermediate to higher protein concentrations, although, we have observed that after a certain higher concentration, the sigmoidal kinetics does not vary much with the total protein concentration significantly. However, our model simulation performed at very low total protein concentration, reestablish the fact that a minimum concentration is required (> 15 *μ*M) to maintain the sigmoidal nature of amyloid formation kinetics ([19]) (**Figure S3**).

### Model accounts for the various time scales associated with the aggregation kinetics in a concentration dependent manner

In spite of being optimized with the quantitative kinetic data related to a specific concentration, our model captures the various features of the kinetics of concentration dependent amyloid formation adequately. To understand this issue further, we further quantified few important time constants such as *τ*_*lag*_ (the time required for the accumulation of the nucleus) and *τ*_50_ (the time required for the amyloid formation reaction to reach 50% completion), normally associated with the kinetics (schematically shown in **Fig. 3A**) for different initial total protein concentrations.

**Fig. 3.**
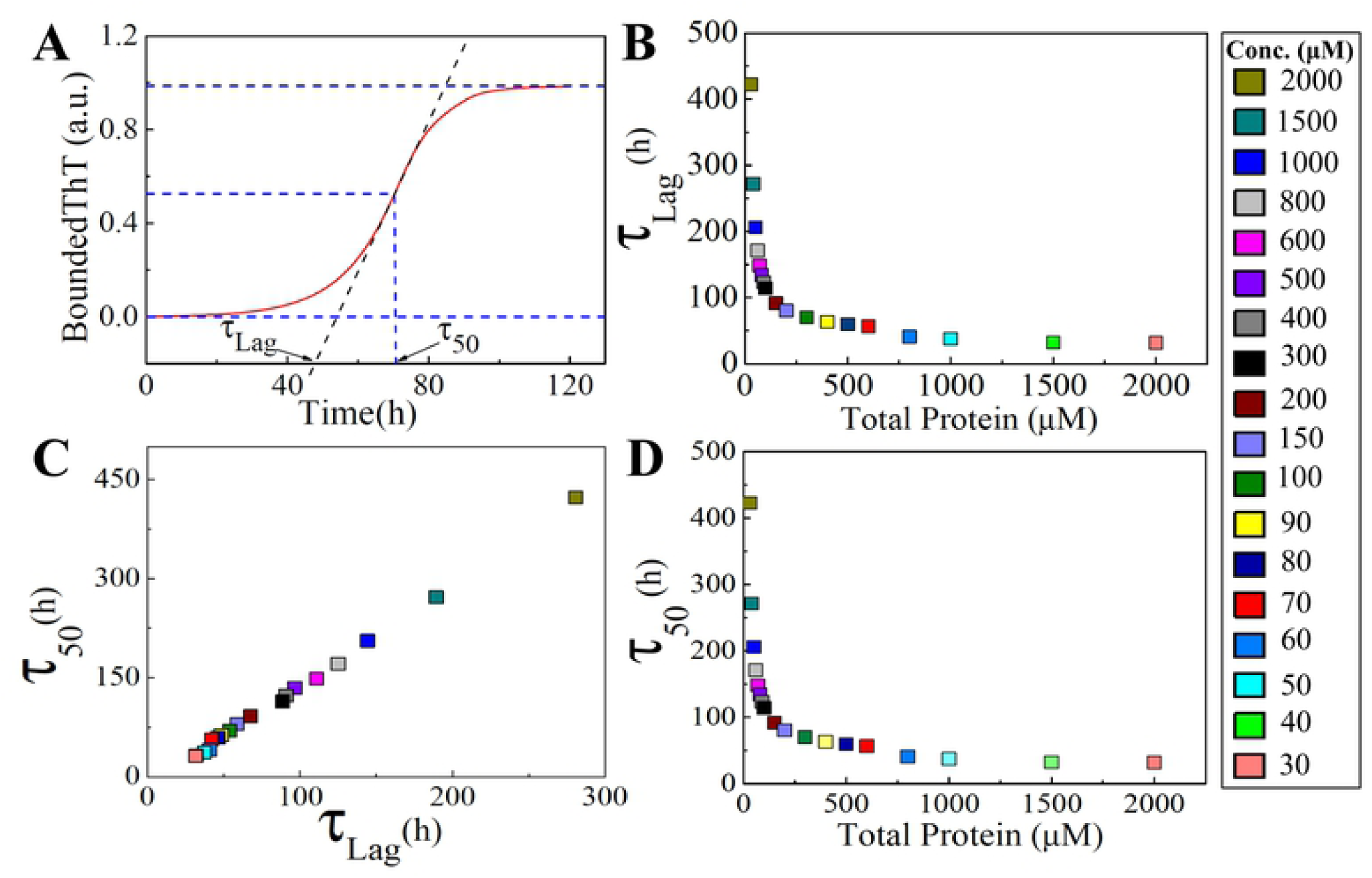
Model accounts for the timescales *τ*_*lag*_ and *τ*_50_ in a concentration dependent manner. (**A)** Schematic definition of the time duration of the lag phase (*τ*_*lag*_) and the duration of the time taken to reach half of the saturation level of bounded ThT (*τ*_50_) at any given total protein concentration. **(B)** A plot of variation of *τ*_*lag*_ as a function of total protein concentration calculated from the model predicted simulated time-courses. Plot of *τ*_*lag*_ with respect to total protein concentration describes at lower and intermediate concentration, *τ*_*lag*_ is linearly correlated where as at higher concentration *τ*_*lag*_ is reaching saturation value. **(C)** Model predicted variation of *τ*_50_ as a function of total protein concentration showed similar behavior as observed with *τ*_*lag*_. **(D)** A correlation plot of *τ*_*lag*_ Vs. *τ*_50_ shows high degree of linear correlation at any given total protein concentration.

We computationally measure how the *τ*_*lag*_ (**Fig. 3B**) and *τ*_50_ (**Fig. 3D**) vary as a function of total protein concentration. From the observation, three distinct regime of total protein concentration (low, medium and, high) can be easily identified, and the characteristic property of these two time-scales can be evaluated separately in those regimes to manifest the underlying nature of protein aggregation dynamics. At lower concentrations regime (up to 200μM), both *τ*_*lag*_ and *τ*_50_ decreases quite sharply with the increase in the total protein concentration that gets reflected in the widely spaced time profiles of the bounded ThT (**Fig. 2C**). However, for intermediate (between 200-600 μM) concentration, the quantitative value of *τ*_*lag*_ and *τ*_50_ does not vary much resulting the bounded ThT curves being closely spaced, but still the fibril formation kinetics can be predicted within the purview of classical nucleation dependent polymerization concept. This gets further confirmed by observing the fact that both the *τ*_*lag*_ and *τ*_50_ decreases quite sharply with the increase in the total protein concentration in a highly correlated manner (**Fig. 3C**). This highly correlated characteristic reveals the behavior of *τ*_*lag*_ and *τ*_50_ as expected for a classical nucleation dependent polymerization reaction.

It turns out that the parameters set estimated by our optimization process, quantified both these time scales with reasonable precision. Due to this, the changes that appear in the initial lag and the elongation phase durations, as a function of the total protein concentrations, are correctly represented in the numerically predicted kinetics (**Fig. 2C**). Only for higher concentrations (> 600μM), the profiles for both *τ*_*lag*_ and *τ*_50_ do not vary significantly, and become independent of total protein concentration indicating the failure of the requirement of classically nucleated polymerizations. One of the probable explanations for this deviated nature may be at higher concentration all the reaction fluxes behave irreversibly as the monomeric concentration of the protein become very high.

### Dynamics of the basic elongation unit has an important role to dictates the pattern of concentration dependent amyloidogenesis

Intriguingly, it has been speculated experimentally that the intermediate protein aggregates (defined as basic elongation units in our model) may be a remarkable kinetic controller of the fibrillation process[12]. We want to elucidate this analysis further by asking whether the duration of the basic elongation units are highly associated with the time scales *τ*_*lag*_ and *τ*_50_. Does the dynamics of these basic elongation aggregates play any important role in influencing the kinetics of amyloidogenesis during the transitions from either lag phase to elongation phase or elongation phase to plateau phase? To inspect this issue, we measured the time scale *τ*_*DI*_ (duration of basic elongation unit, **Fig. 4A**) for different total protein concentration, which was introduced and precisely quantified by Ghosh et al. through experiments ([12]).

**Fig. 4.**
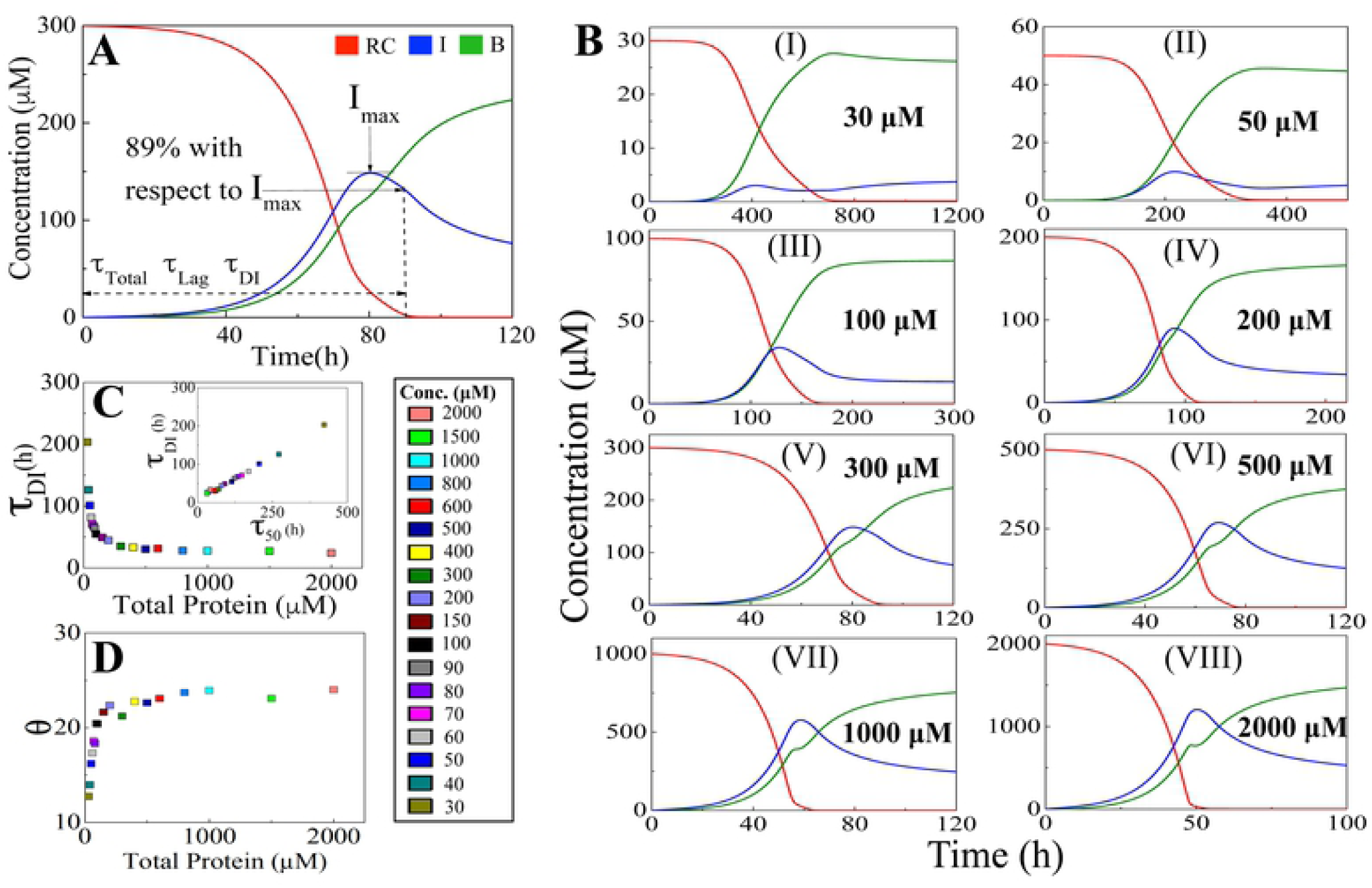
Unraveling the dynamical influence of basic elongation aggregates on amyloid forming kinetics. **(A)** Simulated trajectories of RC, I and B from the best-fitted model for 300 *μ*M total protein concentration, where RC, I and B stands for the total random coil state, total basic elongation aggregates and total fibril like aggregates, respectively. The timescale corresponding to the duration of intermediate (*τ*_*DI*_) measured from experimental observation made by Ghosh et al. corresponds to 89% drop of the I_max_ in case of total protein concentration 300 *μ*M ([12]). **(B)** Model predicted time profiles of RC (Red line), I (Blue line) and B (Green line) under different total protein concentrations corresponding to (i) 30 *μ*M, (ii) 50 *μ*M, (iii) 100 *μ*M, (iv) 200 *μ*M, (v) 300 *μ*M, (vi) 500 *μ*M, (vii) 1000 *μ*M and (viii) 2000 *μ*M, respectively. **(C)** A plot of *τ*_*DI*_ as a function of total protein concentration, where we observe a higher degree of correlation under various total protein concentrations between *τ*_*DI*_ and *τ*_50_ (**inset**). **(D)** A plot of *θ* (in %) against total protein concentration illustrates the effect of basic elongation aggregates in shaping up the amyloid forming kinetics.

In their work, experimentally they have quantified the lag time (*τ*_*lag*_) and the duration of basic elongation unit (*τ*_*DI*_) for 300 *μ*M total protein concentration, where the estimated duration of the sum of these two timescales corresponds to the 89% drop with respect to the I_max_ (**Fig. 4A**). In our case, I_max_ is the maximum level of the total basic elongation aggregates obtained by simulation that correspond to the intermediate aggregates in the experiment performed by Ghosh *et al*. ([12]) We used this knowledge consistently to quantify the *τ*_*DI*_ from our model predicted time profiles (**Fig. 4B**) of total elongation aggregate concentration for different total protein levels. Our analysis reveals that *τ*_*DI*_ varies with total protein concentration (**Fig. 4C**) in a similar fashion like *τ*_*lag*_ and *τ*_50_. Moreover, there is a higher degree of correlation existing between *τ*_*DI*_ and *τ*_50_ (**Fig. 4C**, inset) suggesting that the dynamics of basic elongation aggregates do influence the amyloid forming kinetics in a significant manner.

To better quantify how and at what extent the basic elongation aggregates control the kinetic transition from elongation phase to plateau phase as a function of total protein concentration, we further defined a relative time scale in *θ* (mathematically expressed as 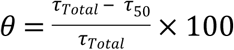, where *τ*_*Total*_ = *τ*_*Lag*_ + *τ*_*DI*_, as defined in **Fig. 4A**). *θ* delineates the influence that the basic elongation aggregates have over the amyloid kinetics after the bounded ThT levels essentially crossed the time points demarcated by *τ*_50_ for different total protein concentration (**Fig. 2C**). In this regard, considering a relative time scale like *θ* makes more sense, as changing the total protein concentration essentially alters all the associated time scales such as *τ*_*Lag*_, *τ*_50_ and *τ*_*DI*_ appreciably (**Fig. 4B**). Our analysis reveals that the value of *θ* systematically increases as a function of total protein concentration, and around 500 *μ*M of total protein concentration, it settles down to a steady level (**Fig. 4D**). This evidently demonstrates that the basic elongation aggregates start influencing the transition from elongation to the plateau phase progressively with the increase in the total protein concentration. Such an effect eventually saturates once the total protein concentration crosses a certain threshold level.

### Model reveals the amyloid formation kinetics get modulated through seeding experiments

The analysis performed in the previous section demonstrates the significant role played by the intermediate aggregate (represented as the basic elongation unit) in modulating the aggregation kinetics. In this context, it is worth mentioning that in the mechanism following the nucleation dependent polymerization, experimentally, the slow rate-determining nucleation step can be evaded by adding the preformed fibril or existing nuclei. This is achieved by either reducing or eliminating the characteristic lag time associated with the amyloidogenesis ([13],[8],[18],[37],[38], which enormously accelerates the amyloid formation. At this point, we further investigate the approach towards numerically performed seeding experiments as similar to *in vitro* experiments by adding a subtle amount of either basic elongation unit or fibril like aggregates at the beginning of the numerical operation. **Fig. 5A** illustrates the investigation that the kinetics of amyloid formation become considerably faster resulting of a much shorter lag time compared to the absence of the seeding experiment (green line). This model predicted analysis qualitatively corroborates with the experimental findings, and suggests that both basic elongation unit and fibril-like aggregates have the capacity in the advancement of different intermolecular reactions, which finally assist to fasten the entire process of amyloidogenesis. Importantly, our model simulations, for the first time exhibit how these basic elongation unit have the ability to influence the lag time and subsequently the amyloid kinetics significantly in a modeling setup.

**Fig. 5.**
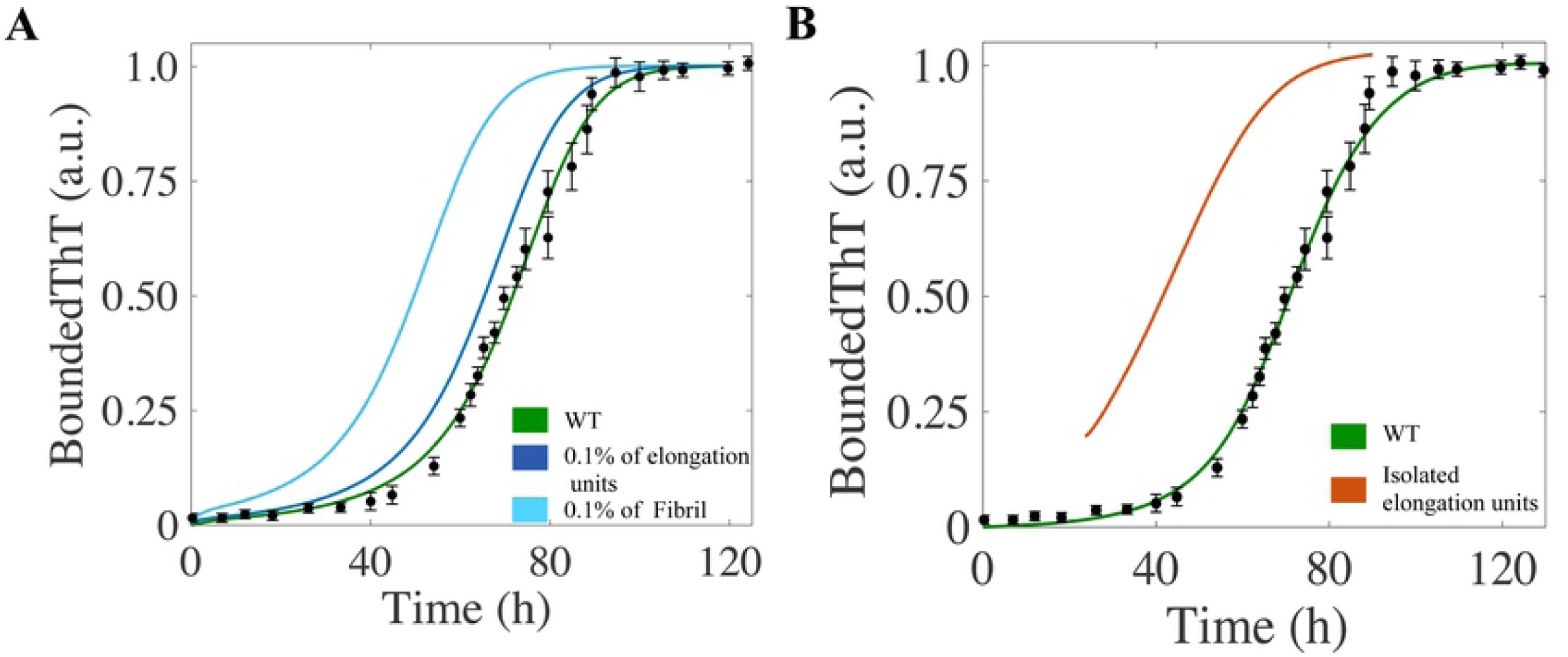
Model reconciles the amyloid formation kinetics of seeding experiments. **(A)** Enhanced rate of the formation of the amyloid fibrils in presence of initial seed added either in the form of basic elongation unit like aggregates (blue line, I_1_=0.1) or fibril like aggregates (magenta line, B_1_=0.1) in comparison to the control simulation performed for the WT (green line) without any seeds. In all the cases, the total protein concentration is 300 *μ*M, and parameters used are taken from **Table S4. (B)** Model simulation with only isolated basic elongation unit aggregates (orange line) predicts that the kinetics of amyloid fibril formation of these aggregates will have no lag time in comparison to experiment performed with monomeric protein (green line). The numerical simulation (orange line) is performed by taking total protein concentration as 200 *μ*M (almost similar in amount of basic elongation unit aggregates isolated experimentally by Ghosh et al. [12]and initial conditions are set up as I_4_=10, M_4_=5, AC=5.

Additionally, experiments further displayed that once isolated, intermediate aggregate can produce amyloid fibril even without going through any lag phase[12]. We have implemented a similar numerical experiment by assuming that we have isolated the basic elongation unit aggregate, which represents the excrementally isolated intermediate aggregate and performed numerical operation for the time evaluation of the fibril formation. The simulation results **Fig. 5B** exhibits that in the absence of the random coil like structure, the only tractable elongation unit have the capability to directly convert into the fibril like aggregates totally hopping the lag phase. One of the very remarkable key features from this observation is that the existence of the random coil like state for fibrillation is not essential, the elongation unit or the pre-fibrillar species i.e. any kind of secondary structure has the capability to form amyloid fibril like structure.

### Model explains the aggregation dynamics of point mutated IDP’s from a dynamical perspective

Now, we challenged the potential of our proposed kinetic model in the regards of explaining the various dynamics of single-point mutated any amyloid forming protein (here we have taken α-Synuclein protein), which seems extremely difficult to achieve in our modeling framework as our model does not contain any kind of protein sequence. However, point mutations in amyloid forming proteins (such as α-Synuclein) often display very interesting dynamical aspect accelerating or slowing down the rate of amyloidogenesis in comparison to the wild type case and lead to different disease states ([39],[40],[41],[42],[43]).

For example, in case of mutant variants of *α*-Synuclein, commonly known as A53V and A53T tend to accelerate the amyloid formation kinetics ([44],[45],[46]), whereas, mutants such as A53K, A53E and A30P is known to behave in a completely opposite manner, elongating the amyloid formation kinetics ([47],[45],[48],[46]). The mutants such as A53V and A53T show a greater tendency to form fibril like aggregate, but have a reduced propensity to form the intermediate elongation like units due to structural changes induced by the respective point mutations ([47]). On the contrary, the other set of mutants (A53K, A53E and A30P) reduce both the propensities to form intermediate and fibril like aggregate. In our modeling framework, we deal with such kind of structural information’s from a dynamical perspective by relatively altering the corresponding reaction rate associated with such different individual reaction mechanism compared to the WT protein. Comparing the relative tendency of the particular mutant and adjusting the specific reaction rate, the model reproduces the kinetics behavior in case of both faster **(Fig. 6A)** and slower mutants (**Fig. 6B**), which qualitatively agrees with the experimental observation. This investigation leads to a remarkable intimation that if the relative fundamental property of any particular amino acid at different stages of fibrillization is well known then the aggregation propensity of any type of mutant will be predictive from our proposed model. This may be helpful in the context of monitoring the kinetics of any mutant as the process is time consuming and there are many complications and challenges from the experimental approaches.

**Fig. 6.**
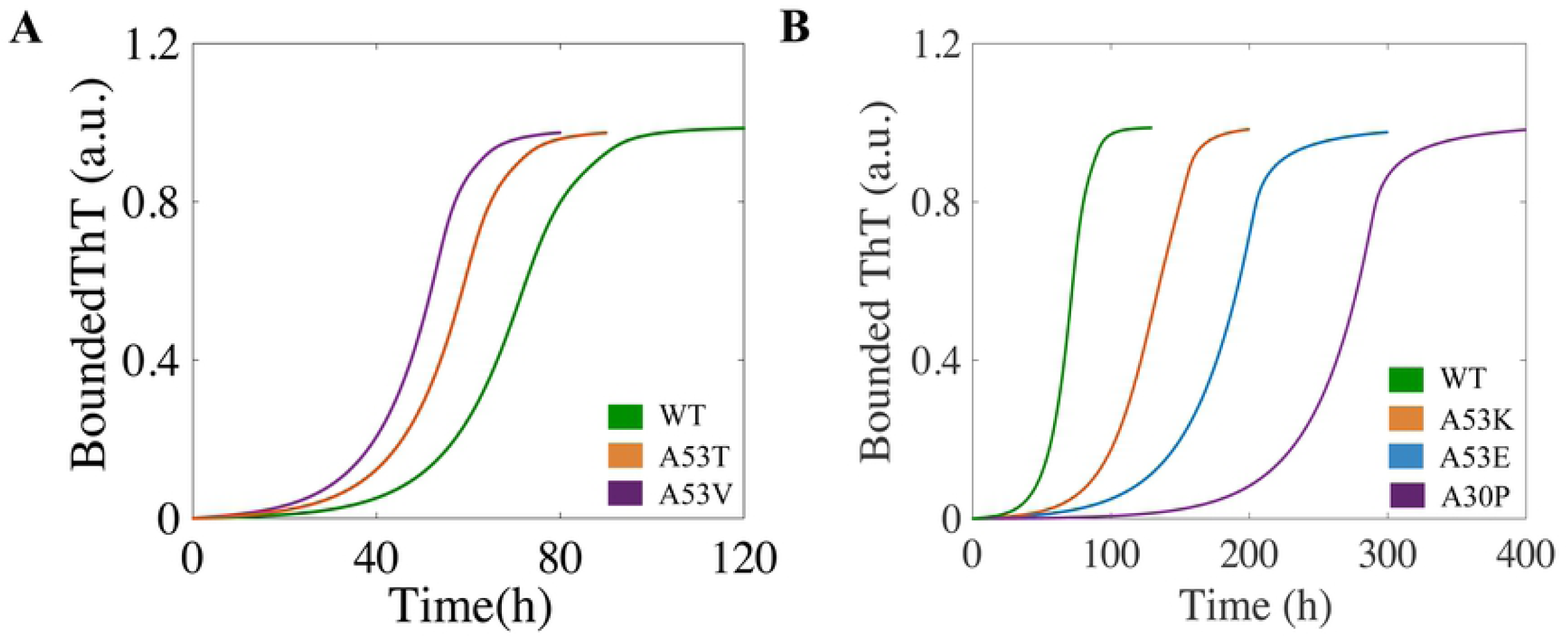
Model explains the aggregation dynamics of point mutated IDP’s. Dynamics of different mutants obtained from our predicted model. The green line represents the WT bounded ThT profiles for both the cases keeping the rate constant as **Table S4. (A)** The time courses obtained for the point mutated IDP’s, the faster mutant like A53T and A53V, by performing simulation separately keeping the rate constant as (*K*_*s*_ = 0.08, *K*_*e*_ = 4.2 for A53T) and (*K*_*s*_ = 0.08, *K*_*e*_ = 4.2 for A53V). **(B)** The simulated trajectory obtained for the point mutated IDP’s, the slower mutant like A53K, A53E and A30P, by performing simulation separately keeping the rate as (*K*_*s*_ = 0.08, *K*_*e*_ = 0.5, *K*_*c*_ = 3.0 for A53K), (*K*_*s*_ = 0.07, *K*_*e*_ = 0.04, *K*_*c*_ = 2.5 for A53E), and (*K*_*s*_ = 0.01, *K*_*e*_ = 0.005, *K*_*c*_ = 0.1 for A30P). (All other parameters are taken as given in **Table S4** for the WT)

### Model makes experimentally tractable predictions to fine-tune the dynamics of amyloidogenesis

Till now, we have shown that our model qualitatively reproduces several features of aggregation mechanism in a generic way. Can our model make experimentally feasible predictions to alter this aggregation process? We tried to answer this question in a methodical way. We employed the sensitivity analysis (**Figure S4**) of the kinetic rate constants (**Table S4**) for the WT case by taking various sensitivity criteria (like *τ*_*lag*_, *τ*_50_ and the slope at *τ*_50_). A careful analysis of **Figure S4** uncovers that some of these kinetic rate constants related to specific steps of aggregation dynamics can prove to be decisive to modify the amyloid forming kinetics in an appreciable fashion. First, we focused on the primary nucleation step, and systematically varied the associated forward rate constant of primary nucleation (K_*x*_) to either inhibit or activate the primary nucleation process computationally. We observe that decreasing K_*x*_ (i.e., inhibiting primary nucleation) leads to significant increase in the lag time without changing the other features of the usual sigmoidal kinetics (**Fig. 7A**) found in case of WT. Interestingly, similar observation has been made for Amyloid-beta 42 (A*β*42) protein in presence of a specific molecular chaperone DNAJB6, where a low sub-stoichiometric ratio of the chaperone produces identical kinetics (as observed in **Fig. 7A**) by inhibiting the primary nucleation event ([49]).

**Fig. 7.**
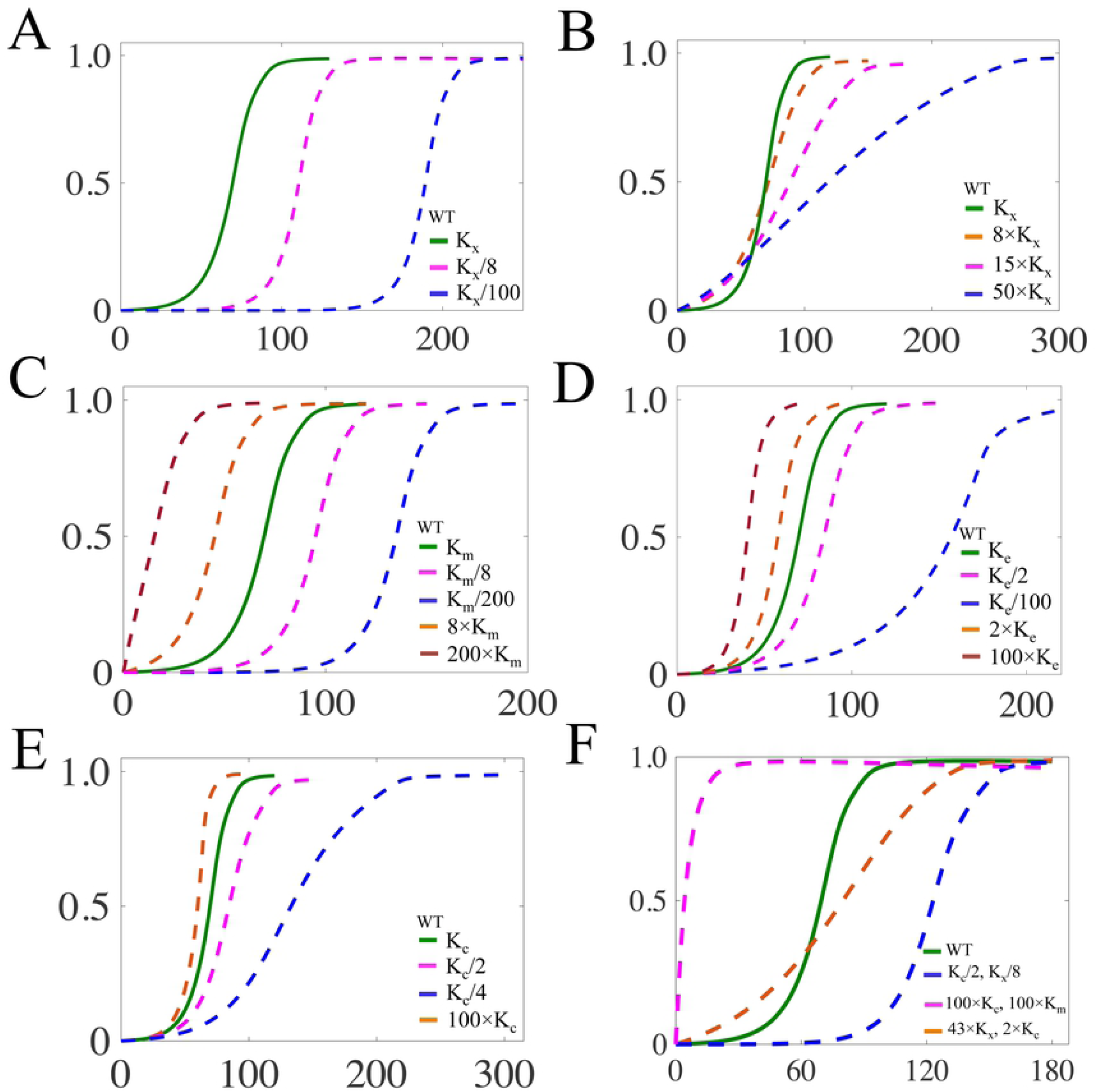
Model predicts ways to fine-tune the aggregation dynamics by perturbing the network responsible for amyloidogenesis. Time courses of bounded ThT profile by perturbing different individual microscopic mechanism obtained by numerical operation through changing with consequent reaction rate constant. Affecting the amyloid forming kinetics by only **(A)** decreasing and **(B)** increasing the rate of primary nucleation at different extent (in comparison to the WT case). Altering the dynamics of amyloid formation by individually influencing the **(C)** conformation transition rate (K_*m*_), **(D)** Transition rate (K_*e*_) of the basic minimum fibril unit (B_1_) from the highest ordered elongation unit, and **(E)** Elongation rate of the fibrillar units (K_*c*_). (**F**) Model predicted kinetics of amyloid formation, when two different rates in amyloid forming network are simultaneously perturbed (either increased or decreased) up to a different degree. In all the figures the changes in the kinetics are relative to the WT case (300 *μ*M protein concentration, green line) and the parameters used are given in **Table S4**.

The model further predicts that an increase in K_*x*_ (i.e., facilitating the primary nucleation) can produce two contrasting scenarios. While a little increase in K_*x*_ reduces the lag time and preserves the other features of sigmoidal growth kinetics (**Figure S5A**), whereas a moderate to higher values of K_*x*_, significantly changes the nature of aggregation kinetics (**Fig. 7B**). This implies that at higher rate of primary nucleation, the lag time will be reduced, but the other process associated to fibrillization becomes relatively slower. To better understand this anomaly, we looked at the dynamics of various species involved in the process (**Figure S5B-C**). **Figure S5** unravels that at moderate to higher nucleation rate, the amount of monomer (M_1_) very quickly converts into higher order aggregate (without any substantial conformational transition, M_4_). Thus, monomeric species (M_1_) are consumed such that they are not available for the elongation steps to form the higher order elongation units and fibril forming units elongating these process which has been depicted in **Fig. 7A**. This prediction can be verified by activating the nucleation process in a specific manner with the help of experimental technique.

Thereafter, we concentrated on the two most important steps of the aggregation kinetics; (i) the transformation process of the highest order random coil state (M_4_) into activated state (AC) and the corresponding kinetic parameters K_*m*_, and (ii) the transition from a higher order elongation unit (I_4_) to the basic minimum fibril forming unit (B_1_) and the related kinetic parameter K_*e*_. Subsequent analysis predicts that increasing either K_*m*_ or K_*s*_ (transition rate of Activated complex (AC) to form basic minimum elongation unit I_1_) eventually shortens the lag time and *τ*_50_ with comparably very little raises in the slope at *τ*_50_ (**Fig. 7C, Figure S6A)**, which inflected from the sensitivity analysis studies (**Figure S4)**.

However, a decreasing value of K_*e*_ not only increases the lag and *τ*_50_ (**Fig. 7C**), but also creates significant changes in the slope of the aggregation kinetics at *τ*_50_ as speculated by the sensitivity graphs (**Figure S4**).

Finally, we dealt with the most crucial step of amyloid forming kinetics, which comprises of the fibril elongation and the secondary nucleation events. In our model, these two steps are controlled by the two kinetic rate constants K_*c*_ and n_1_. Our model simulations predict that even a significant increase in the fibril formation rate (K_*c*_) will marginally affect the quantities like *τ*_*lag*_, *τ*_50_ and the slope at *τ*_50_ in comparison to the WT situation, whereas decreasing K_*c*_ does not effect the *τ*_50_ much but appreciably modifies the *τ*_50_ quantity, specially generating enormous changes in the slope at *τ*_50_ (**Fig. 7E**). We found similar outcome, when we specifically varied the parameter n_1_ to see the effect of secondary nucleation process separately (**Figure S6B**). This indicates that methodically modifying the fibril elongation process in one way or other, one can affect the amyloid formation kinetics in a significant way. Importantly, our model predicted aggregation kinetics (**Fig. 7E** or **Figure S6B**) has a striking qualitative resemblance with the experimental findings, where the secondary nucleation events are inhibited by using molecular chaperones belonging to Brichos family ([50]). These results (**Fig. 7A-E** or **Figure S5-7**) evidently portray the potential of our predicted model to qualitatively forecast the dynamics of amyloid aggregation, when a distinct microscopic intermolecular reaction mechanism related to aggregation process is perturbed.

### Model predicts rational drug designing strategies to control dynamics of amyloid formation

Next we asked whether we can obtain any further insight about amyloid forming kinetics, if we simultaneously target more than one specific steps associated with the amyloidogenesis? This is an important question as nowadays one can experimentally design different kinds of molecular chaperones or specific molecular inhibitors to control amyloid kinetics by concurrently targeting different steps of aggregation process. However, rationalizing the molecular basis of these experimental observations in a precise way is extremely challenging. To this end, we performed model simulations by varying rates of nucleation, basic elongation unit formation and secondary nucleation in pairs and in different extent to gain further perception in this context. We found increasing both primary nucleation rate (K_*x*_) and the transition rate (either K_*m*_ or K_*s*_) of M_4_ to the basic minimum elongation unit (I_1_) facilitates the amyloid formation immensely by drastically reducing the *τ*_*lag*_ and *τ*_50_ (**Fig. 7F**, pink dashed line). This result is in quite stark contrast, when only K_*x*_ is significantly elevated (**Fig. 7B**), but resemble more with the situation, where either of the transformation rates (K_*m*_ or K_*s*_) were increased appreciably (**Fig. 7C** or **Figure S6A**). This emphasizes the fact that the transformation step from M_4_ to I_1_ is an extremely crucial step for the aggregation process. A moderate increase in primary nucleation rate (K_*x*_) supplemented with a small increase in the secondary elongation rate (K_*c*_) does not affect the *τ*_*lag*_ and *τ*_50_ much, but changes the in the slope at *τ*_50_ (**Fig. 7F**, orange dashed line) in comparison to the situation when only (K_*x*_) is varied (**Fig. 7B**). However, partially inhibiting both the primary nucleation and secondary elongation steps will lead to an abrupt increase (**Fig. 7F**, blue dashed line) in *τ*_*lag*_ and *τ*_50_ compared to **Fig. 7A** (pink dashed line), where primary nucleation has been disrupted.

In a similar note, Xu *et al*. had shown that one could use a specific inhibitor (Ssa1p) at different concentrations to inhibit the fibril elongation rate of Ure2p protein, and the aggregation dynamics varied both in terms of lag time and the slope of the aggregation kinetics at the half maxima point in comparison to the WT case ([51]). We can qualitatively reconcile the similar aggregation dynamics by simultaneously altering the related kinetic constants associated with the elongation process. Our model simulations essentially predicts that by simultaneously reducing the rates of transition from activated complex to elongation unit (*K*_*s*_, affects lag phase significantly), the formation rate of initial fibril seed (*K*_*e*_) and the rate corresponding to fibril elongation and secondary nucleation (*K*_*c*_) at different extent, one can qualitatively capture the inhibitory effect induced by molecular chaperone Ssa1p during the kinetics of aggregation of Ure2 protein [51]. Thus our analysis implies that this kind of molecular chaperones affect the kinetics of amyloid formation by simultaneously targeting various steps associated with amyloidogenesis.

These observations suggest that our model simulations can qualitatively elucidate and interpret the effect of various external perturbations made to modify the amyloid aggregation network in a dynamical manner. Additionally, using our model, we can gain important mechanistic insights about the aggregation process in presence of any external perturbations that probe more than one step related to amyloid formation. These insights can have serious implication while designing drug molecules to regulate the process of amyloid formation in an efficient and controlled manner and can lead to novel therapeutic strategies to counter amyloid formation kinetics.

## Conclusion

Protein aggregation process that leads to amyloid formation is undoubtedly a complex phenomenon. Unraveling the detailed mechanism, via which the monomeric proteins transform into an amyloid like fibril state and inflict several disease phenotypes, is extremely challenging to unfold by only using conventional biochemical techniques. Recent experimental advancement to probe molecular aggregation at the single molecule level might be the way forward to disentangle the complicated set of intermediate events that organize the amyloidogenesis, but at the moment these techniques are still at the primitive state. However, to produce effective therapeutic measures to control the amyloid formation demands a better understanding about the underlying dynamical events, which orchestrate amyloid fibril formation. In this work, we explored a generic protein aggregation network (**Fig. 1** and **Figure S1**) by employing chemical kinetic analysis (**Table S2**) with an aim to interpret the aggregation kinetics under different experimental conditions from a dynamical viewpoint. Importantly, we wanted to investigate whether such a model system can be realized, which will be able to qualitatively predict and elucidate the experimental observations related to amyloid formation.

In this regard, we have been able to systematically identify an appropriate generic model out of many variants (**Table S5**) of amyloidogenesis that has been optimized by using relevant experimental kinetic data (**Fig. 2**) to obtain a reliable set of kinetic constants (**Table S4**) for the aggregation network. Our model simulations not only reconcile various time scales (**Fig. 3**) associated with the amyloid formation as the function of initial protein concentrations, but it further provides insight behind such observations (**Fig. 4**) by qualitatively explaining various experimental findings performed with the WT (**Fig. 5**) and mutant variants (**Fig. 6**) of the amyloid forming proteins. This clearly suggests that our model, which considers a relatively detailed protein interaction network of aggregation and developed on the basis of experimental data, can corroborate the dynamics of amyloid formation kinetics efficiently.

However, the success of any mathematical model lies in it’s predictive ability exploiting which we gain further insights about how to probe the concerned network in an optimal way for our advantage. In this context, our model proved to be extremely effective to predict a diverse range of amyloid dynamics (shown schematically in **Fig. 8**) that can emanate, if one strategically influences various parts of the aggregation network either individually or collectively at different extent (**Fig. 7** and **Figure S5-7**). **Fig. 8** evidently demonstrates that a multitude of amyloid forming kinetics having different *τ*_*lag*_, *τ*_50_ and slope at *τ*_50_ can be qualitatively realized by either inhibiting or activating various steps responsible for amyloidogenesis in a systematic and rational way. These predicted aggregation kinetic patterns would be helpful in generating precise small molecule inhibitors (or specific molecular chaperones) to prevent unwanted amyloidogenesis by targeting specific steps in the aggregation network. Additionally, it will also provide us with the detailed mechanistic insight about how the network of aggregation got perturbed under such kind of inhibitory conditions. Moreover, the conclusions drawn from our model simulations (**Fig. 8**) can be explicitly used to fabricate functional amyloid and novel biomaterials.

**Fig. 8.**
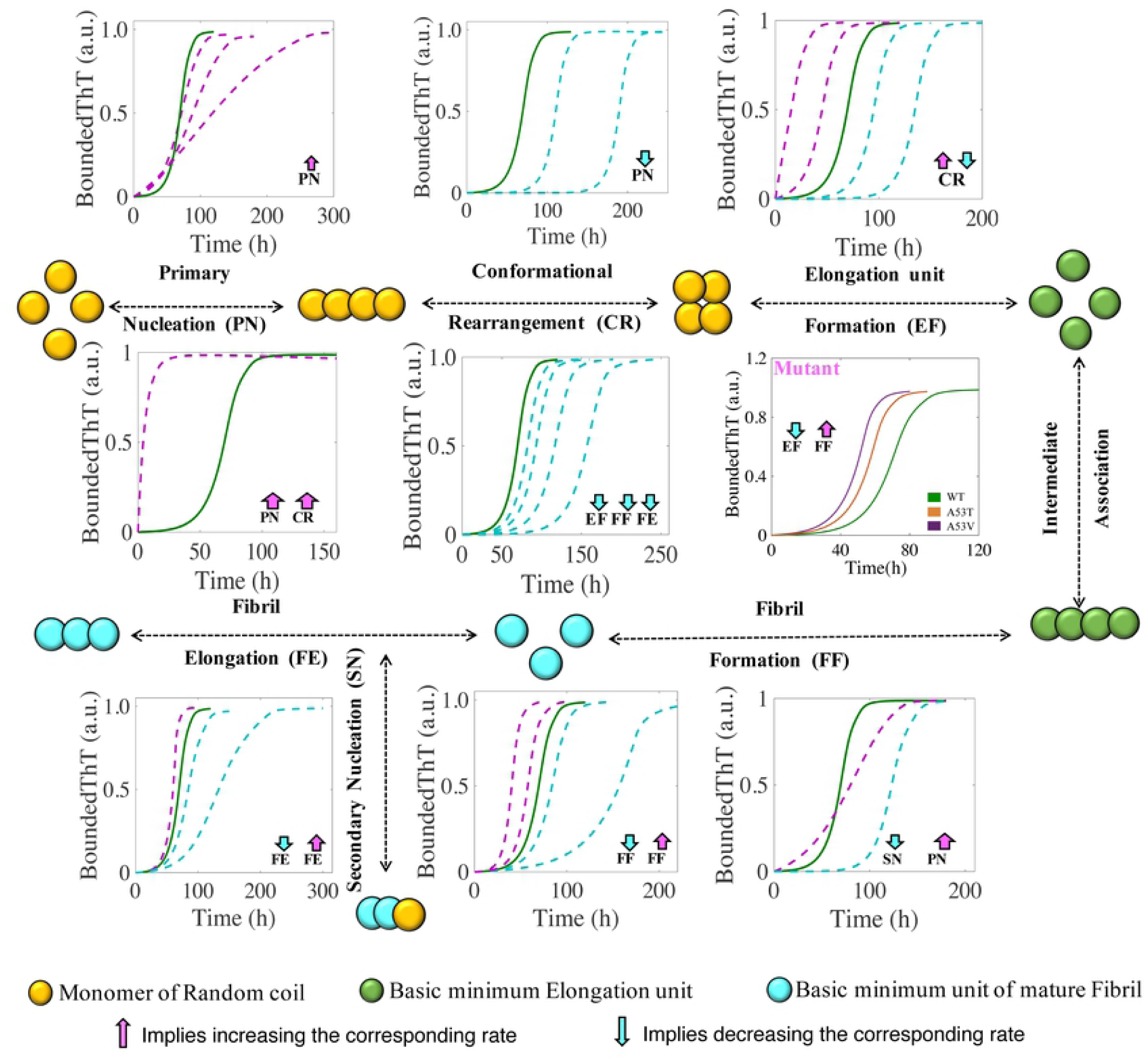
Model demonstrates various possibilities to alter the amyloid forming kinetics by providing a mechanistic insight. A schematic description of the protein aggregation network depicted in **Fig.1**, wherein we showed how perturbing different steps involved in amyloid forming kinetics either individually or collectively can influence the nature of the dynamics of amyloidogenesis. All the kinetics is drawn with respect to the WT case (***Green line***) for total protein concentration 300 *μ*M, where we used the parameter set given in **Table S4**. The corresponding amyloid kinetics obtained by either increasing or decreasing various rates is shown with ***Pink*** and ***Blue lines***, respectively.

In summary, our mathematical modeling strategy provides a generic framework to study the amyloid forming kinetics in a comprehensive manner for the diverse kind of proteins that are prone to form such aggregates in one way or other. Although the modeling approach does not consider the structural information’s of the monomeric protein, it offers an alternative path to understand the effect of point mutations from a dynamical perspective. Importantly, the mechanistic insights behind different kinetic profiles attained by altering the rates at distinct points in the network revealed the possibilities of developing potent inhibitors in the form of drug molecules to resist amyloid formation. We believe that our modeling approach will found wide applicability in coming days to decipher the mechanism of amyloid formation in different context.

## Method

The regulatory network (**Fig.1** or **Figure S1**) was constructed in term of ordinary differential equations (ODE). The kinetic parameters of the model have been estimated using the PottersWheel software (version 4.1.1) ([52]). The parameters were fitted globally in logarithm parameter space employing genetic algorithm method, where the CVODE method is used for integrating the ODE’s. Each model system is optimized (∼500 times) by taking several random initial starting values of the parameters, and several parameter sets were obtained after the optimization process, which fit the experimental data adequately. We have only shown the best fitted results in each case for different models that are considered in Table S5. Each parameter estimation round was started with parameter values that were disturbed with a strength of *ϵ* = 0.2, so that *P*_*New*_ = *P*_*Current*_ × 10^(*ϵ* × *ϕ*)^ with *ϕ* being normally distributed with mean 0 and variance 1. The detail description of the development of entire models and their relative χ^2^ value and Akaike information criterion (AIC) has been provided in **S2 Text**. The χ^2^ value was calculated according to the following equation-

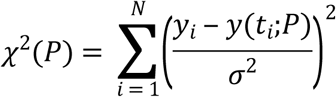

where, *y*_*i*_ being the experimental data points and *y*(*t*_*i*_;*P*) is the simulated value at time point *i* for parameters value *P, σ* is the standard deviation of the experimental data set. Once the best fitted parameter set for any model is obtained after optimization, the rest of the simulation and analysis has been done in freely available software XPPAUT by considering corresponding ODE files for the respective models.

## Acknowledgements

Thanks are due to IIT Bombay for providing fellowship to BT.

---

**Fig S1**: **Detail interaction network for aggregation process of amyloid forming protein**. All the variables corresponding to this network is defined in **Table S1**, the differential equation corresponding to each variable are shown in **Table S2 and S2 Text**, and the kinetic rate constants are defined in **Table S4**. Origins of all the reaction events are explained in **S1 Text**. Here, **X** represents the elongation rate of the intermediate elongation unit, the forward rate constant could be either ***K***_***i*1**_ or ***n*** × ***K***_***i***_ and the backward rate constant could be either ***K***_***j*1**_ or ***n*** × ***K***_***j***_. **Y** represents the elongation rate of the intermediate fibril unit; the forward rate constant could be either ***K***_***c*1**_ or ***n*1** × ***K***_***c***_ and the backward rate constant could be either ***K***_***d*1**_ or ***n*1** × ***K***_***d***_. The pink colour symbol represents the ThT binding with secondary structure of protein.

(TIFF)

**Fig S2**: **The best-fitted time courses of bounded ThT and % of total basic elongation unit for all the different models**. The time course simulation of all different variants (given in Table S5) is shown together. The experimental data set considered for each specific model is same as mentioned in the main text [1]. The model equations are changed according to the particular network configuration considered as mentioned in the **S2 Text**.

(TIFF)

**Fig S3: Numerical analysis of bounded ThT at different low total protein concentrations**. Time course simulation at 20*μM*, 15*μM* and 10*μM* total protein concentrations. Model simulations predict that for 15*μM* and 10*μM* concentrations, although the simulation has been done keeping the scaling factor at its highest value (F=1), the amyloid kinetic profiles do not saturate. This signifies the fact that a minimum concentration (here 15*μM* or below) is needed for the protein to drive the process of amyloidogenesis under the parametric condition provided in **Table S4**.

(TIFF)

**Fig S4**: **Sensitivity analysis of the kinetic rate constants are performed by taking *τ***_***lag***_, ***τ***_**50**_ **& slope of *τ***_**50**_ **as the sensitivity criteria**. Each of the kinetic rate constants are increased about 20% of the given values provided for the WT case (**Table S4**) and the sensitivity is measured therein. This analysis dictates the sensitivity of the rate constant associated with nucleation, elongation seed formation, association, intermediate elongated seed association, fibril seed formation, fibril elongation & secondary nucleation on lag time, *τ*_50_ & slope of *τ*_50_, which characterize the time scale measurement and sensitivity to kinetic parameters evaluated after fitting experimental data of the aggregation process.

(TIFF)

**Fig S5**: **Quantitative dynamical analysis of protein aggregation upon increasing the primary nucleation rate. (A)** Changes in the global kinetic profile of the aggregation process of the amyloid protein. The kinetics of protein aggregation accelerates when the primary nucleation rate (K_x_) is increased to a low extent decreasing the lag phase and subsequently accelerating the overall kinetics of the aggregation process as described in the figure. However, increasing the primary nucleation rate to a greater extent (Fig. 7B) decrease the lag phase but slows down the other process. In **(B)** & **(C)**, we have plotted the time profiles of various forms of the protein aggregates to understand the reason behind this observation. In both these graphs, the solid line represents the kinetic profile of WT situation, and the dotted line represents when the primary nucleation rate (K_x_) has been increased by 50 times. Increase in the primary nucleation rate elevates the amount of M_2_, M_3 &_ M_4_ and reduces the amount of M_1_ drastically **(Fig-S2C)**. As a result, we observe a reduction in lag phase (**Fig-7B**). Since other rate constants were kept constant, the conversion of M_4_ to the next higher order structure taking much time compare to WT. Now, with very reduced amount of Monomer (M_1_), the elongation process of elongation unit and secondary nucleation of fibril gets obstructed which need the presence of monomer and thus it leads the delay in the further processes **(Fig-S2B)**. Thus the overall kinetics after the lag phase becomes very slow that has been reflected in BThT curve in **Fig-7A**.

(TIFF)

**Fig S6**: **Model predicts ways to alter the aggregation dynamics by perturbing different steps in the self-assemble process responsible for amyloidogenesis. (A)** The kinetics of amyloid formation accelerates if the formation of the basic minimum elongation unit (I_1_) becomes faster than the kinetics compared to WT, while reducing the same rate (*K*_*s*_) leads to delay in dynamics of amyloid formation. For both the cases slope at *τ*_50_ remains almost unaltered demonstrating alteration of *K*_*s*_ only affects *τ*_*lag*_. (B) Proliferating the secondary nucleation rate (by varying only *n*1) by increasing the corresponding rate speeded up the aggregation process not much on the lag time, but not much as effective as the kinetics have been effected by K_s_. Instead inhibition of the secondary nucleation process creates significant changes on the sharp rise of the kinetics specially controlling the elongation process. So, varying the secondary nucleation rate alter the sharp growth rather that the lag time, as a result the slope of the *τ*_50_ changes.

(TIFF)

**Fig S7**: **Model simulations qualitatively produce the amyloid forming kinetics that resemble with the aggregation kinetics of Ure2 prion protein in presence of Ssa1p**. The green solid line represents the WT (*K*_*s*_ = 0.145,*K*_*e*_ = 1.411, *K*_*c*_ = 7.503) case, orange line represents the dynamics when *K*_*s*_ = 0.1,*K*_*e*_ = 1.0, *K*_*c*_ = 7.2, blue line denotes BThT dynamics for *K*_*s*_ = 0.07, *K*_*e*_ = 0.8, *K*_*c*_ = 7.0, purple line denotes the situation for *K*_*s*_ = 0.04,*K*_*e*_ = 0.5, *K*_*c*_ = 6.5, and the sky-blue line represents the sigmoidal dynamics when *K*_*s*_ = 0.01,*K*_*e*_ = 0.35, *K*_*c*_ = 6. In the presence of molecular chaperone Ssa1p, the kinetics of the aggregation of Ure2 protein gets inhibited due to the interaction of Ssap1 with the oligomeric protein that are eventually populated during and after the lag phase like elongation seed or fibril seed [9]. By decreasing the *K*_*s*_, which is the transition rate from activated complex to elongation unit effecting the duration of lag phase significantly, by reducing the *K*_*e*_ which is the formation rate of initial fibril seed, by changing the *K*_*c*_ which effect both the elongation of fibril and secondary nucleation rate, we are getting same kind of kinetic observation from our mathematical model which resemblance with experiment observations. Gradually decreasing in all the rate constant results in delay of the aggregation kinetics, which agreed with the experimental consequences that with increasing the molar ration of Ssap1, the duration of the fibrillation kinetic gets extended.

(TIFF)

**Table S1. Abbreviated name of the species involved in the model**.

**Table S2. Ordinary differential Equations governing the protein aggregation process**.

**Table S3. Algebraic Equations**.

**Table S4. Description of the parameters involved in the Model and Parameter Value for the best-fitted Model-1A**.

**Table-S5. Comparing an ensemble of model configurations to identify a most probable network & a dynamic model of amyloidogenesis by performing statistical analysis**.

**S1 Text. Quantitative dynamic modeling of the aggregation process of amyloid forming Protein**

**S2 Text. Comparison of different model configurations**

